# A critical role of AREG for bleomycin-induced skin fibrosis

**DOI:** 10.1101/2021.01.20.427509

**Authors:** Mary Yinghua Zhang, Shuyi Fang, Hongyu Gao, Xiaoli Zhang, Dongsheng Gu, Yunlong Liu, Jun Wan, Jingwu Xie

## Abstract

We report our discovery of an important player in the development of skin fibrosis, a hallmark of scleroderma. Scleroderma is a fibrotic disease, affecting 70,000 to 150,000 Americans. Fibrosis is a pathological wound healing process that produces an excessive extracellular matrix to interfere with normal organ function. Fibrosis contributes to nearly half of human mortality. Scleroderma has heterogeneous phenotypes, unpredictable outcomes, no validated biomarkers, and no effective treatment. Thus, strategies to slow down scleroderma progression represent an urgent medical need. While a pathological wound healing process like fibrosis leaves scars and weakens organ function, oral mucosa wound healing is a scarless process. After re-analyses of gene expression datasets from oral mucosa wound healing and skin fibrosis, we discovered that several pathways constitutively activated in skin fibrosis are transiently induced during oral mucosa wound healing process, particularly the amphiregulin (*Areg*) gene. *Areg* expression is upregulated ~10 folds 24hrs after oral mucosa wound but reduced to the basal level 3 days later. During bleomycin-induced skin fibrosis, a commonly used mouse model for skin fibrosis, *Areg* is up-regulated throughout the fibrogenesis and is associated with elevated cell proliferation in the dermis. To demonstrate the role of Areg for skin fibrosis, we used mice with *Areg* knockout, and found that *Areg* deficiency essentially prevents bleomycin-induced skin fibrosis. We further determined that bleomycin-induced cell proliferation in the dermis was not observed in the *Areg* null mice. Furthermore, we found that inhibiting MEK, a downstream signaling effector of Areg, by selumetinib also effectively blocked bleomycin-based skin fibrosis model. Based on these results, we concluded that the Areg-EGFR-MEK signaling axis is critical for skin fibrosis development. Blocking this signaling axis may be effective in treating scleroderma.

## BACKGROUND

Scleroderma affects 70,000 to 150,000 Americans^1,2^. Due to the heterogeneous phenotypes, unpredictable outcomes, no validated biomarkers, and no effective treatment, scleroderma is a very hard disease to manage in the clinic. Similar to other fibrotic diseases, scleroderma is a pathological wound healing process that produces an excessive extracellular matrix to interfere with normal organ function^3–6^. Fibrosis is known to contribute to nearly half of human mortality^7^. While molecular mechanisms underlying fibrosis remain elusive, fibrosis shares several commons processes. Chronic injury in tissues is the initial trigger for fibrosis, which draws inflammatory responses^3^. Effector cells, such as fibroblasts, become activated in order to repair tissue damages. During the repair process, extracellular matrix (ECM) is produced. Unlike scarless wound repair in oral mucosa ^8^, fibrosis is caused by excessive ECM production from over-activated effector cells. As a result, fibrotic tissues are formed, leading to abnormal tissue function or even organ failure.

One critical step in fibrosis is production of excessive ECM from over activated effector cells. In contrast to fibrosis, oral mucosa wound healing has less ECM and controlled number of effector cells^9^. We hypothesize that by comparing scarless oral mucosa wound healing with skin fibrosis on gene expression, we may find molecules that are critical for scleroderma development^8^. Since there are many studies on the time course of gene expression during oral mucosa wound healing and skin fibrosis, we took the advantage of public datasets to re-analyze gene expression during oral mucosa wound healing. Specifically, we are interested in those genes with transient induction after injury in oral mucosa^9^. These genes will be further analyzed for their expression at different time points during skin fibrosis.

We found that *Areg* and the ErbB signaling pathway is transiently induced during oral mucosa wound healing. In contrast, *Areg* is highly expressed throughout the process of skin fibrosis. Using *Areg* knockout mice and a specific inhibitor of downstream effector MEK, selumetinib, we demonstrated that the Areg-EGFR-MEK signaling axis is critical for bleomycin-induced skin fibrosis.

## RESULTS

### Gene expression comparison between oral wound and skin fibrosis

We hypothesized that the genes, that are transiently induced during oral mucosa wound healing but upregulated during skin fibrosis, may be critical for driving skin fibrosis. Several public datasets are available on gene expression during oral cavity wound healing, skin wound and skin fibrosis. We chose to use GSE23006 in which gene expression data from multiple time points during oral wound healing are available ^9,10^. GSE132869 (from Geo datasets) was used for re-analyses of gene expression during bleomycin-induced skin fibrosis.

In analyses of GSE23006, we focused on the genes induced 12hr and 24hrs after wound but returned to the basal level (+/−20%) 3 days after injury. Only genes induced over 2 folds after injury were further analyzed for their expression in bleomycin-mediated skin fibrosis. As shown in ***Fig.S1***, 91 genes were induced shortly after injury but returned to the basal level 3 days later. Gene ontology analysis indicates that two of these genes are ErbB-related molecules AREG and HBEGF (***Fig.S1, Fig.1A***). Others include molecules related to immunity, keratinization, antivirus defense and chemotaxis (***Fig.1A***).

**Figure 1.**
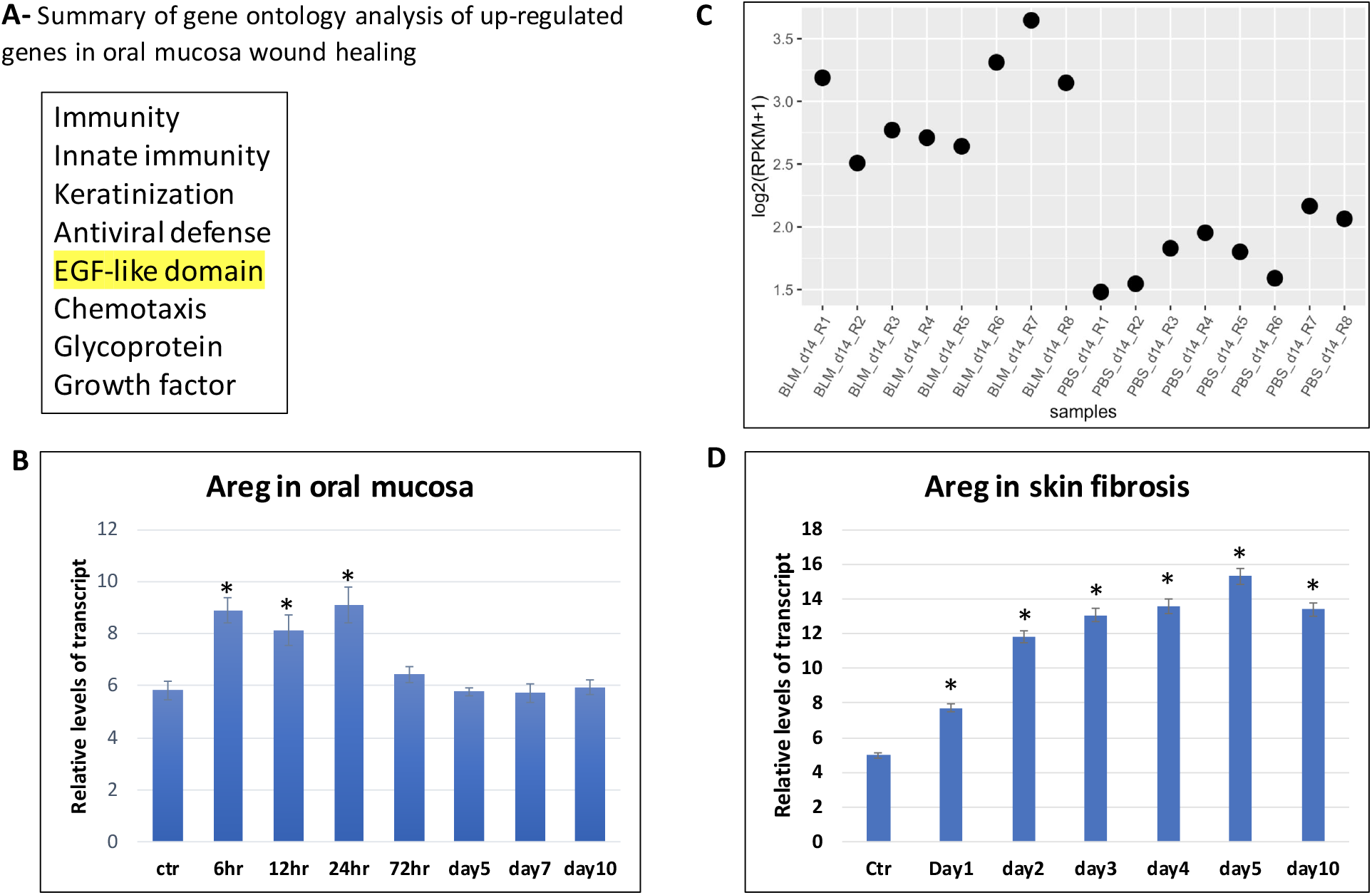
Areg expression in oral wound healing and skin fibrosis. **A** shows a summary of ontology analysis of transient up-regulated genes during oral mucosa wound healing. The list of genes with transient induction is shown in Supplementary Table 1. **B** shows a re-analysis of Areg expression during oral mucosa wound healing ^8^, and * indicates a significant change (p<0.01). **C** shows re-analysis of Areg gene expression from GSE132869 (from Geo datasets), confirming the significant expression in bleomycin-induced skin fibrosis. Each dot represents one sample. PBS was used as a control, and BLM indicates bleomycin treatment. **D** shows time-dependent expression of Areg during bleomycin-induced skin fibrosis. Please note that Areg was consistently highly expressed in the skin biopsies after bleomycin treatment at different time points. * indicates a significant change (p<0.01).

We further analyzed expression of these genes in GSE132869, and 9 genes were up-regulated in bleomycin-induced skin fibrosis (***Fig.S2***). These molecules include molecules known to be involved in fibrosis, such as *Mmp9, Cxcl1, Ccl5, Fosl1* and *Stat1*. For example, Mmp9 is known to be important for skin fibrosis ^11–13^. The function of other 3 genes, *Klra2, Slfn4* and *Hdc*, during fibrosis is not known. In contrast, the pattern of *Areg* expression is very intriguing. *Areg* was induced to ~10 folds shortly after oral cavity wound, returned to the basal level 3 days later (***Fig.1B***), but was up-regulated throughout bleomycin-induced skin fibrosis (***Fig.S2, Fig.1C***). We have confirmed elevated *Areg* expression during skin fibrosis using skin specimens at different time points (***Fig.1D***). Since Areg is a known growth factor with well-characterized signaling downstream effectors (Areg-EGFR-RAS-RAF-MEK), we focused our efforts to determine the role of Areg signaling for bleomycin-mediated skin fibrosis.

### Skin fibrosis in Areg null mice

Because *Areg* is one of the few genes with transient induction during oral wound healing but maintains high expression throughout skin fibrosis, we used *Areg* knockout mice to determine the significance of *Areg* for bleomycin-induced fibrosis.

We used skin thickness as a way to measure skin fibrosis. Skin thickening is one of the earliest manifestations of scleroderma; and is one of the most widely used measures in the clinical trials ^14–17^. Several studies have demonstrated that the extent of skin involvement directly correlates with internal organ involvement and prognosis in scleroderma patients.

We obtained *Areg* knockout mice and wild type mice to perform bleomycin-induced skin fibrosis through daily intradermal injection of bleomycin (50 microliter of 1mg/ml bleomycin in PBS) for 10 days. At the end of the study, skin biopsies at the injected site or at a site far away the injection site were harvested to perform H&E staining, and the skin thickness, which is the combination of epidermis and dermis, was measured by Image J. In wild type mice, bleomycin injection increased the skin thickness ~80-150% (***Fig.2***, P <0.001). In *Areg* null mice, however, we only observed <20% increase of skin thickness after intradermal injection of bleomycin. The data from gene knockout mice demonstrates that *Areg* is critical for skin fibrosis.

**Figure 2.**
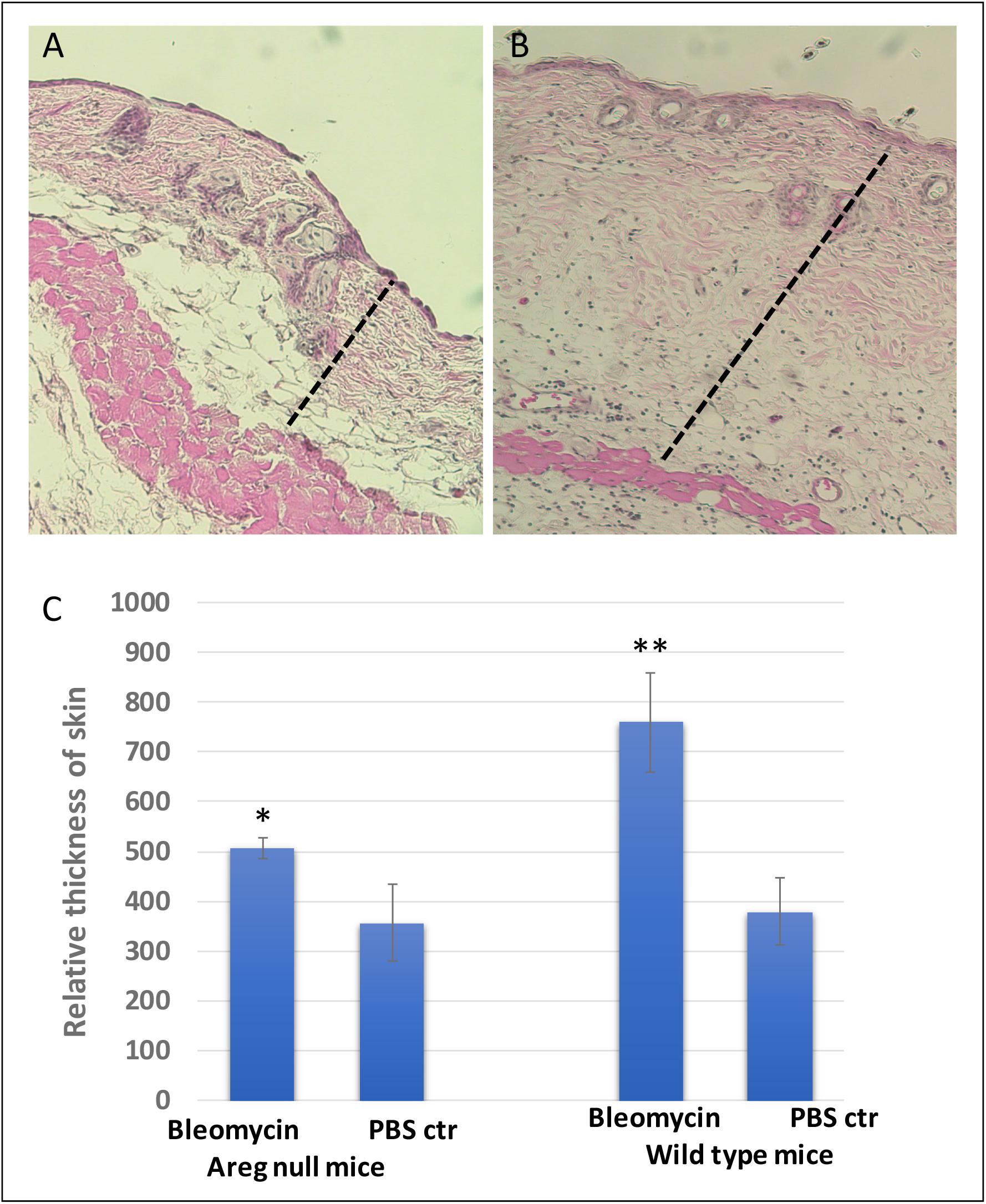
A significant role of Areg in bleomycin-induced skin fibrosis. **A** & **B** show H&E images of skin biopsies from control mice treated with PBS control (A) or bleomycin (B). **C** shows a summary of data from 10 mice for each group (5 females and 5 males from wild type or *Areg* knockout). * indicates < 0.05, and ** indicates p<0.0001. Dashed bars indicate skin thickness.

Increased proliferation of fibroblasts and other cell types is accompanied with fibrosis, and the level of cell proliferation has been used to monitor disease progression or treatment outcomes in the mouse models^18–20^. We detected expression of Ki-67 in skin specimens following bleomycin injection to determine the specific effects on cell proliferation and fibrosis progression in mice. We detected an increase in Ki-67 positivity in dermis of bleomycin-injected mice (***Fig.3***), indicating an elevated cell proliferation during skin fibrosis. Our data are consistent with previous studies to show elevated cell proliferation during fibrosis^21^. In *Areg* null mice, however, bleomycin did not increase the level of Ki-67 positivity in skin dermis, suggesting a role of Areg for cell proliferation in skin dermis during fibrosis (***Fig.3***).

**Figure 3.**
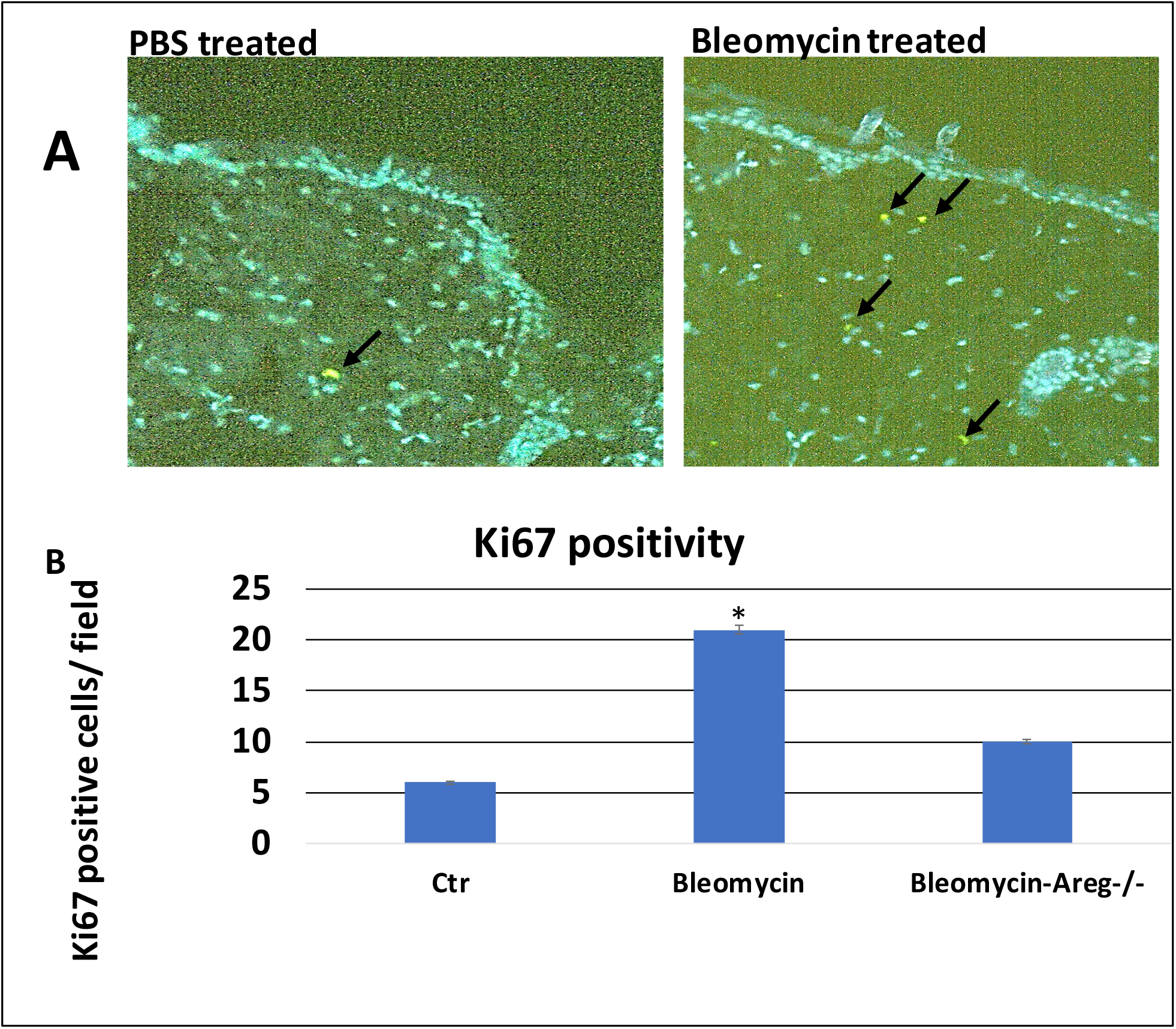
Ki67 positivity in skin biopsies. We detected Ki67 positivity in skin biopsies either treated with PBS or bleomycin through immunofluorescence (IF) staining with ki67 specific antibodies. **A** shows images with positive Ki67 staining (green, indicated by arrows). **B** shows a summary of ki67 staining from 6 mice (3 females and 3 males) in each group. * indicates p < 0.0001.

### Blocking MEK signaling prevents bleomycin-induced skin fibrosis

There are several downstream effectors of Areg, including the receptor EGFR, MEK and PI3K. Selumetinib (also AZD6244) is a specific MEK inhibitor, now approved by FDA to treat children 2 years and older with neurofibromatosis type I ^22–24^. We hypothesized that if MEK mediates Areg effects in skin fibrosis, treatment of selumetinib should reduce bleomycin-mediated skin fibrosis.

First, we determined that MEK is activated in skin fibrosis. Using specific antibodies to phospho-ERK, we found that a high percentage of phospho-ERK positive cells in the skin from mice treated with bleomycin (***Fig.4***). Next, we treated mice with selumetinib (oral gavage, 15mg/kg/every other day), together with intradermal injection of bleomycin (50 microL of 1mg/ml bleomycin in PBS) for 2 weeks and determined skin thickness after H&E staining of the skin biopsies. As shown in ***Fig.5***, we found that while wild type control skin increased its thickness over 80% after bleomycin treatment, selumetinib treated skin tissues had <10% of increase in skin thickness. This result demonstrates that selumetinib is effective in reducing bleomycin-induced skin fibrosis.

**Figure 4.**
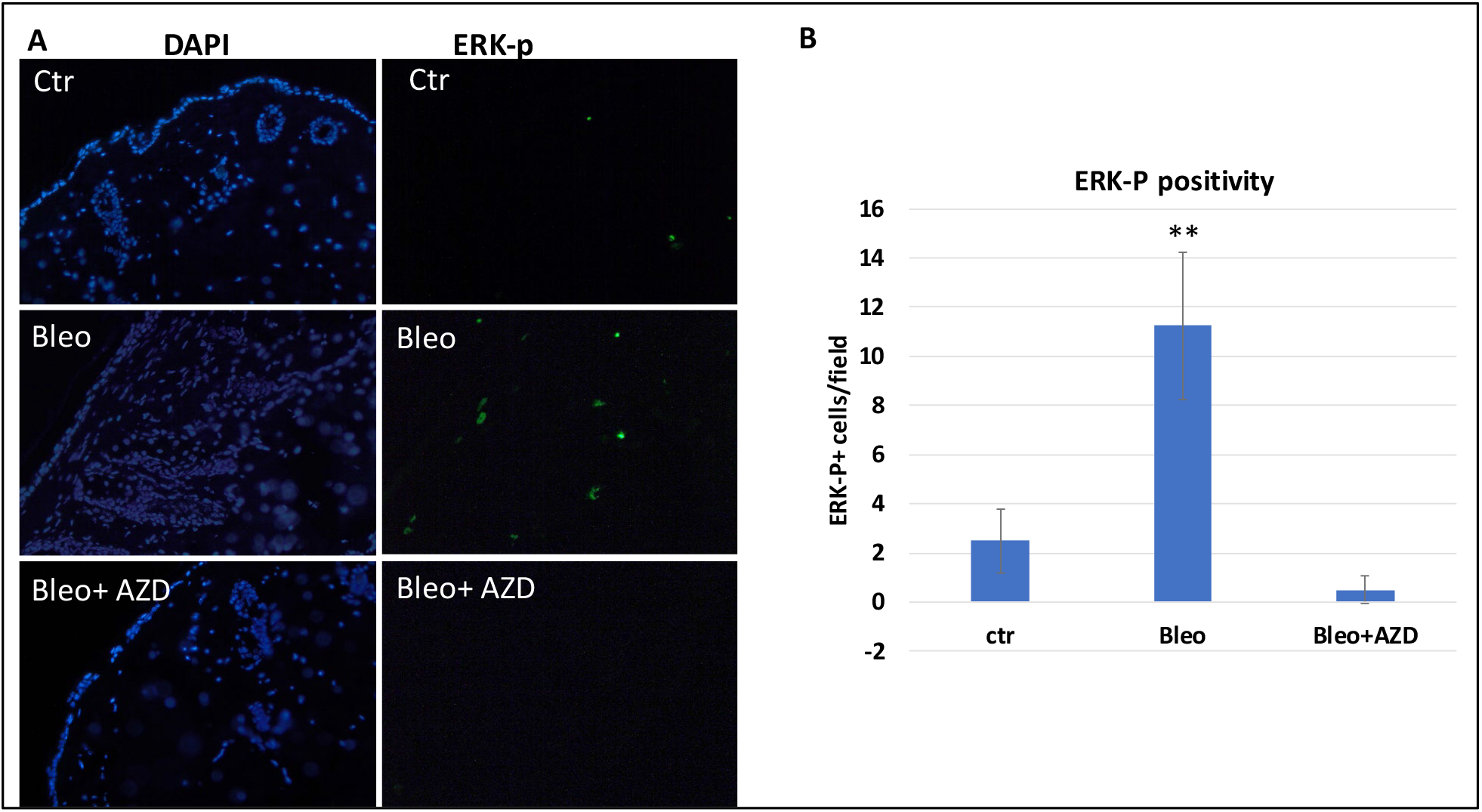
ERK-p expression in skin fibrosis and after AZD6244 (selumetinib) treatment. **A** shows images of nuclear staining with DAPI or ERK-p specific antibodies. The method is described in details in experimental methods. **B** is a summary of ERK-p staining using the number of positive cells per field under microscope (200 X). ** indicates p <0.0001.

**Figure 5.**
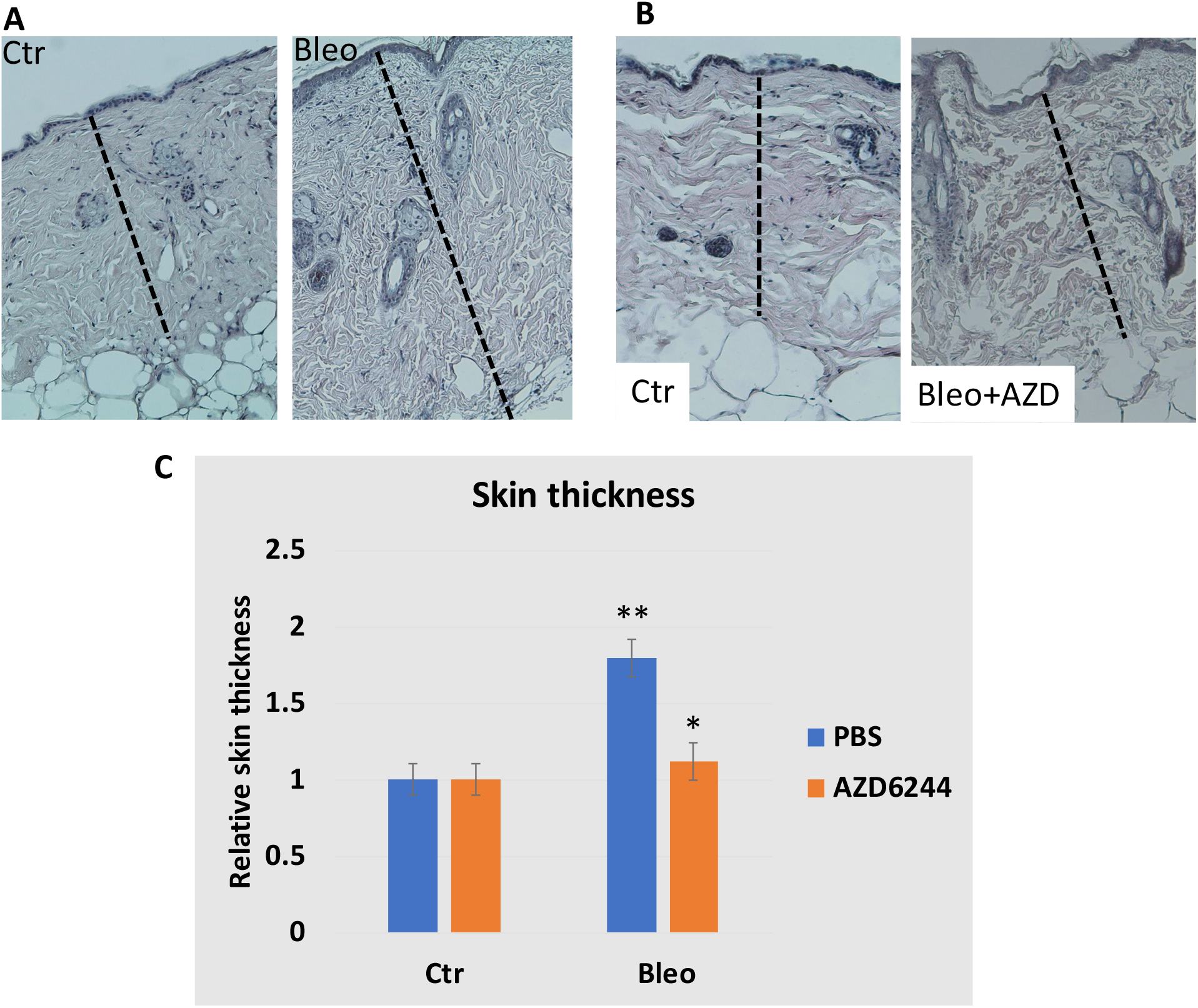
The effect of MEK inhibitor AZD6244 (shown as AZD, also selumetinib) on bleomycin-induced skin fibrosis. **A** shows typical H&E images of skin biopsies from bleomycin-based mouse model of skin fibrosis. **B** shows H&E images from mice treated with AZD6244 during bleomycn-induced skin fibrosis. **C** shows the summary of the data from two groups of mice treated with AZD6244 or PBS (n=3). Dashed bars indicate skin thickness. ** indicates p <0.0001, and * indicates p= 0.045.

We further examined the level of phospho-ERK in selumetinib-treated skin specimens and found that phospho-ERK is undetectable after selumetinib treatment (***Fig.4***).

Our data using selumetinib further demonstrate that the Arge-EGFR-MEK signaling axis is critical for development of skin fibrosis.

## DISCUSSION

Fibrosis is associated with changes in many cell types and a variety of signaling events, many of them interconnected. There are several approaches to dissect the signaling networks critical for the development of fibrosis: 1) analyses of human specimens to determine the signaling changes in fibrosis^25–27^; 2) use of mouse models of fibrosis to dissect changes at different times during fibrosis^28^; 3) comparing different wound healing processes to identify the abnormalities in the fibrosis process^9^. Our approach is to compare the differences between oral mucosa wound healing and skin fibrosis with the assumption that fibrosis is a pathological wound healing process. Thus, identifying the abnormalities in the fibrotic process may help establish new ways to restore the tissue remodeling process. Indeed, we have proved that Areg, a molecule transiently induced in oral wound healing but constitutively upregulated in skin fibrosis, is critical for development of bleomycin-induced skin fibrosis, a hallmark of scleroderma. We further showed that inhibiting downstream signaling of Areg by MEK inhibitor selumetinib is as effective as Areg knockout in prevention of skin fibrosis development. Based on these data, we predict that inhibitors of the Areg-EGFR-MEK signaling axis may be effective in treatment of scleroderma and other fibrotic diseases.

We showed the significance of Areg for skin fibrosis, but the exact mechanisms of action remain to be determined. Using single cell RNA-seq, we generated data to indicate that Areg expresses in at least two cell populations: immune cells and fibroblasts (or myofibroblasts) (***Fig. S3***). Further analyses of Areg signaling in these two cell populations will help lineate the signaling network of Areg during skin fibrosis. Previous studies showed that Areg is highly expressed in type 2 innate lymphoid cells (ILC2), which is induced by IL3 3^29^. It is also reported that Areg can be regulated by TGFbeta signaling or Hippo-YAP signaling ^30–32^. Our single cell RNA-seq showed correlative expression of IL33 and Areg in immune cells, suggesting a role of IL33 in regulation of Areg. Additional analyses of downstream effectors may also help understand Areg signaling during skin fibrosis. Our data with MEK inhibitor selumetinib not only showed the significance of the Areg-EGFR-MEK signaling axis for fibrosis, but also provided novel ways to treat/prevent fibrosis. Whether blocking the Areg-EGFR-MEK signaling axis is effective in treatment of skin fibrosis will require additional experiment using mice with established skin fibrosis.

Are our data relevant to human scleroderma? Gene expression of bulk scleroderma specimens of skin often does not reveal up-regulation of *AREG* in the diseased biopsies although a recent report showed up-regulation in scleroderma patients ^33^. However, activated immune cells from scleroderma show a high level of Areg expression^34^, indicating that expression of *AREG* is associated with specific cell types in the disease process, rather than the overall gene expression. Indeed, single cell analyses indicate that *AREG* is highly expressed in myofibroblasts, not other types of fibroblasts in scleroderma patients ^25^. Additional work is needed to show the relevance of Areg signaling in other types of scleroderma specimens, such as lung and kidney specimens at the single cell level.

During our studies, we also identified several other molecules with similar features as Areg, and their roles in fibrosis have not completed proven. While *Fosl1, Mmp9, Cxcl1, Ccl5* and *Stat1* have been linked to fibrosis in the literature, their direct evidence for mechanisms of action during fibrosis remain elusive. On the other hand, there are limited resources for studies of *Slfn4, Klra2* and *Hdc*. One approach will be to use gene knockout mice. Knockout mice of *Mmp9, Cxcl1, Ccl5, Hdc* and Stat1 are viable and may be obtained commercially from Jackson laboratory. Alternatively, neutralizing antibodies to Cxcl1 and Ccl5 can be used to block the function of the targets^35,36^. These studies likely generate additional ways to mitigate scleroderma development and progression.

## CONCLUSIONS

Based on our results, we conclude that Areg and the Areg-EGFR-MEK signaling axis plays an important role in the development of skin fibrosis in bleomycin-based mouse model. Since skin fibrosis is a hallmark of scleroderma, we believe that inhibitors of Areg signaling may be effective in prevention and treatment of scleroderma.

## METHODS

### 1. Chemicals

AZD6244 (selumetinib) was purchased as a research agent from the Selleckchem Chemicals LLC (Houston, TX, USA). AZD6244 (selumetinib), originally developed by *AstraZeneca*, is a highly potent and selective non-ATP competitive inhibitor of MEK with an IC_50_ of 14 nM^37^. Bleomycin Sulfate, Streptomyces verticillus, was purchased from Sigma (St. Luis, MO).

### 2. Animals, bleomycin-based skin fibrosis, and treatment

*Areg* knockout and C57Bl/6 mice were purchased from Jackson laboratory (Bar Harbor, ME). Use of animal was approved by the IACUC committee in Indiana University School of Medicine (ethical code 11370; approval date—15 February 2020). We mated *Areg* knockout and C57Bl/6 mice over 7 generations to obtain *Areg* knockout and wild type mice in a similar genetic background. Genotyping of mice was performed by PCR with specific primers provided by the vendors using lysed tail from each mouse [0.3 cm tail in 100 mL of PCR Direct (tail) solution (Viagen Inc., Los Angeles, CA) with 1 mg/mL proteinase K at 55°C overnight, then 85°C for 45 minutes, and use 0.5–1 microL of the lysate for each 25 microL PCR reaction].

Skin fibrosis was generated according to a previously published protocol ^38^. In brief, bleomycin was made in PBS in 1mg/ml, and intradermal injection was performed using gauge needle #27 with 50microL/injection onto the back of mouse skin (at low right side) after fur removal. At up left, we used PBS for injection after fur removal. The procedure was approved by the IUCUC Committee in Indiana University School of Medicine and was strictly followed in the study. We injected mice daily for 10 days before harvesting skin tissues for analyses in skin thickness, gene expression and histology [hematoxylin and eosin (H&E) staining and Immunofluorescence (IF)]. In bleomycin-induced skin fibrosis studies, we used 10 mice per group (control PBS injection or bleomycin injection, 5 females and 5 males). We used 3 mice for AZD6244 (selumetinib) treatment, and 5 mice for the control group.

AZD6244 was suspended in sterile PBS by sonication at 5 mg/mL. For drug treatment, mice were treated with AZD6244 (oral gavage 10 mg/kg daily) or vehicle control (PBS) in each group.

### 3. Histology, IF staining and tissue measurement

Histology was performed according to a previously published procedure^39^. Fresh tissue was harvested and fixed with 10% buffered-formalin or zinc-based fixative. Five-micron paraffin-embedded sections were labeled with primary antibodies against phosphor-p44/p42 MAPK (Erk1/2) (Thr 202/Tyr 204) (Cell Signaling Technology Cat# 4370, 1:200, Danvers, MA, USA), Ki-67 (ab15580, 1:500, AbCam, Cambridge, MA, USA).

The skin thickness was quantified using ImageJ. To avoid discrepancy from age and genetic backgrounds of the mice, we used littermates from the same mating cage for selection of treatment groups or genotypes. Because of the variation between back skin and abdomen skin in tumor development, we used back skin for injection and histology studies.

### 4. RNA Extraction, RT-PCR and Real-Time PCR

Total RNAs from tissues were extracted using Tri-RNA reagent from Sigma (St Luis, MO) according to the manufacturer’s instruction and 1 microG of total RNA was reverse transcribed into cDNAs using the first-strand synthesis kit (Roche, Tucson, AZ, USA). Real-time quantitative PCR analyses were performed according to a previously published procedure ^39^. Triplicate CT values were analyzed in Microsoft Excel using the comparative Ct(DDCt) method as described by the manufacturer (Applied Biosystems, Foster City, CA, USA). The amount of target (ddCt) was obtained by normalization to an endogenous reference (*Gapdh* for mice) and relative to a calibrator. All TaqMan primers and probes were purchased from Applied Bio systems Inc.

### 5. Single cell RNA-seq, Sequence Alignment, Differential Expression Analyses

Tissue was dissociated with Collagenase IV 1mg/ml 37C for 1 hour. Cells were washed with 5%FBS/PBS twice, each centrifuged (1000rpm), and dead cells were removed by dead cell removal kit (Miltenyi Biotec, Somerville, MA). Prepared cells were used to perform 10X Genomics according to a previously published protocol^40^. In brief, appropriate number of cells were loaded on a multiple-channel micro-fluidics chip of the Chromium Single Cell Instrument (10x Genomics) with a targeted cell recovery of 9,000. Single cell gel beads in emulsion containing barcoded oligonucleotides and reverse transcriptase reagents were generated with the v3.1 Next GEM Single Cell 3□ reagent kit (10X Genomics). Following cell capture and cell lysis, cDNA was synthesized and amplified. Illumina sequencing library was then prepared with the amplified cDNA. The resulting library was sequenced using a custom program on Illumina NovaSeq 6000. 28 bp of cell barcode and UMI sequences and 91 bp RNA reads were generated with Illumina NovaSeq 6000 at CMG of Indiana University School of Medicine. Sequencing analyses were performed as previously reported ^41^.

### 6. Statistical analyses Data

are presented as mean SD. Statistical analyses were performed using the Mann–Whitney test or the Student t test (two-tailed) to compare the results, with P values of <0.05 as statistically significant.

## Supporting information

Gene lists

Figure on single cell RNA-Seq

## LIST OF ABBREVIATIONS

AREG: amphiregulin
ECM: extracellular matrix
MEK: MAPK/ERK kinase
ERK: extracellular receptor-stimulated kinase
MAPK: mitogen-activated protein kinase.
EGFR: Epidermal Growth Factor Receptor
H&E: Hematoxylin and Eosin
ILC2: group 2 innate lymphoid cells
IL33: Interleukin 33

## DECLARATIONS

### ETHICS APPROVAL AND CONSENT TO PARTICIPATE

Use of animal was approved by the IACUC committee in Indiana University School of Medicine (ethical code 11370; approval date—15 February 2020).

### CONSENT TO PUBLICATION

Not applicable

### AVAILABILITY OF DATA AND MATERIALS

Not applicable

### AUTHOR CONTRIBUTIONS

Conceptualization and Design: JX.

Resources: JX.

Methodology: MZ, SF, XZ, GD.

Data Analysis and Interpretation: HG, JW, YL, JX.

Writing, review, and/or revision of the manuscript: JX.

Supervision: JX.

## ACKNOWLEDGEMENT

We thank The Well Center for Pediatric Research, Jeff Gordon Research Laboratory, AGA, Healthcare Initiatives, Inc. and CTSI Indiana for support. Single cell analysis was carried out in the Center for Medical Genomics at Indiana University School of Medicine, which is partially supported by the Indiana University Grand Challenges Precision Health Initiative. This work was supported by the Riley Foundation for Children, AGA and Indiana University School of Medicine Biomedical Research Grant.

## AUTHORS’ INFORMATION

**Mary Yinghua Zhang, Xiaoli Zhang, Dongsheng Gu, Jingwu Xie**

The Wells Center for Pediatric Research, Department of Pediatrics, Indiana University School of Medicine, Indiana;

**Shuyi Fang, Yunlong Liu, Jun Wan**

Department of BioHealth Informatics, Indiana University School of Informatics and Computing at IUPUI

**Hongyu Gao, Yunlong Liu, Jun Wan, Jingwu Xie**

The IU Simon Comprehensive Cancer Center, Indiana University, Indiana

**Yunlong Liu, Jun Wan**

The Center for Computational Biology and Bioinformatics, Indiana University School of Medicine, Indiana;

**Yunlong Liu, Jun Wan**

Department of Medical and Molecular Genetics, Indiana University School of Medicine, Indiana.

**Correspondence should be addressed to Jingwu Xie, Ph.D.,**

The Wells Center for Pediatric Research, Department of Pediatrics, Indiana University School of Medicine, Room R4-327, 1044 W Walnut St., Indiana.

## REFERENCES

1. Ingegnoli F, Ughi N, Mihai C. Update on the epidemiology, risk factors, and disease outcomes of systemic sclerosis. Best Pract Res Clin Rheumatol 2018;32:223–40.

2. Barnes J, Mayes MD. Epidemiology of systemic sclerosis: incidence, prevalence, survival, risk factors, malignancy, and environmental triggers. Curr Opin Rheumatol 2012;24:165–70.

3. Distler JHW, Gyorfi AH, Ramanujam M, Whitfield ML, Konigshoff M, Lafyatis R. Shared and distinct mechanisms of fibrosis. Nat Rev Rheumatol 2019;15:705–30.

4. Nanthakumar CB, Hatley RJ, Lemma S, Gauldie J, Marshall RP, Macdonald SJ. Dissecting fibrosis: therapeutic insights from the small-molecule toolbox. Nat Rev Drug Discov 2015;14:693–720.

5. Rockey DC, Bell PD, Hill JA. Fibrosis--A Common Pathway to Organ Injury and Failure. N Engl J Med 2015;373:96.

6. Wynn TA. Cellular and molecular mechanisms of fibrosis. J Pathol 2008;214:199–210.

7. Wynn TA. Fibrotic disease and the T(H)1/T(H)2 paradigm. Nat Rev Immunol 2004;4:583–94.

8. Glim JE, van Egmond M, Niessen FB, Everts V, Beelen RH. Detrimental dermal wound healing: what can we learn from the oral mucosa? Wound Repair Regen 2013;21:648–60.

9. Chen L, Arbieva ZH, Guo S, Marucha PT, Mustoe TA, DiPietro LA. Positional differences in the wound transcriptome of skin and oral mucosa. BMC Genomics 2010;11:471.

10. Leonardo TR, Shi J, Chen D, Trivedi HM, Chen L. Differential Expression and Function of Bicellular Tight Junctions in Skin and Oral Wound Healing. Int J Mol Sci 2020;21.

11. Vafashoar F, Mousavizadeh K, Poormoghim H, et al. Gelatinases Increase in Bleomycin-induced Systemic Sclerosis Mouse Model. Iran J Allergy Asthma Immunol 2019;18:182–9.

12. Kim WU, Min SY, Cho ML, et al. Elevated matrix metalloproteinase-9 in patients with systemic sclerosis. Arthritis Res Ther 2005;7:R71–9.

13. Waszczykowska A, Podgorski M, Waszczykowski M, Gerlicz-Kowalczuk Z, Jurowski P. Matrix Metalloproteinases MMP-2 and MMP-9, Their Inhibitors TIMP-1 and TIMP-2, Vascular Endothelial Growth Factor and sVEGFR-2 as Predictive Markers of Ischemic Retinopathy in Patients with Systemic Sclerosis-Case Series Report. Int J Mol Sci 2020;21.

14. Matsuda KM, Yoshizaki A, Kuzumi A, et al. Skin thickness score as a surrogate marker of organ involvements in systemic sclerosis: a retrospective observational study. Arthritis Res Ther 2019;21:129.

15. Amjadi S, Maranian P, Furst DE, et al. Course of the modified Rodnan skin thickness score in systemic sclerosis clinical trials: analysis of three large multicenter, doubleblind, randomized controlled trials. Arthritis Rheum 2009;60:2490–8.

16. Clements PJ, Lachenbruch PA, Ng SC, Simmons M, Sterz M, Furst DE. Skin score. A semiquantitative measure of cutaneous involvement that improves prediction of prognosis in systemic sclerosis. Arthritis Rheum 1990;33:1256–63.

17. Zheng B, Nevskaya T, Baxter CA, et al. Changes in skin score in early diffuse cutaneous systemic sclerosis are associated with changes in global disease severity. Rheumatology (Oxford) 2020;59:398–406.

18. He Y, Tsou PS, Khanna D, Sawalha AH. Methyl-CpG-binding protein 2 mediates antifibrotic effects in scleroderma fibroblasts. Ann Rheum Dis 2018;77:1208–18.

19. Mikamo M, Kitagawa K, Sakai S, et al. Inhibiting Skp2 E3 Ligase Suppresses Bleomycin-Induced Pulmonary Fibrosis. Int J Mol Sci 2018;19.

20. Fan Y, Zhang W, Wei H, Sun R, Tian Z, Chen Y. Hepatic NK cells attenuate fibrosis progression of non-alcoholic steatohepatitis in dependent of CXCL10-mediated recruitment. Liver Int 2020;40:598–608.

21. Kendall RT, Feghali-Bostwick CA. Fibroblasts in fibrosis: novel roles and mediators. Front Pharmacol 2014;5:123.

22. Bergqvist C, Wolkenstein P. MEK inhibitors in RASopathies. Curr Opin Oncol 2020; Publish Ahead of Print.

23. Killock D. Selumetinib benefits children with inoperable plexiform neurofibromas. Nat Rev Clin Oncol 2020;17:273.

24. Gross AM, Wolters PL, Dombi E, et al. Selumetinib in Children with Inoperable Plexiform Neurofibromas. N Engl J Med 2020;382:1430–42.

25. Valenzi E, Bulik M, Tabib T, et al. Single-cell analysis reveals fibroblast heterogeneity and myofibroblasts in systemic sclerosis-associated interstitial lung disease. Ann Rheum Dis 2019;78:1379–87.

26. Lu T, Klein KO, Colmegna I, Lora M, Greenwood CMT, Hudson M. Whole-genome bisulfite sequencing in systemic sclerosis provides novel targets to understand disease pathogenesis. BMC Med Genomics 2019;12:144.

27. Apostolidis SA, Stifano G, Tabib T, et al. Single Cell RNA Sequencing Identifies HSPG2 and APLNR as Markers of Endothelial Cell Injury in Systemic Sclerosis Skin. Front Immunol 2018;9:2191.

28. Yue X, Yu X, Petersen F, Riemekasten G. Recent advances in mouse models for systemic sclerosis. Autoimmun Rev 2018;17:1225–34.

29. Cao Q, Wang Y, Niu Z, et al. Potentiating Tissue-Resident Type 2 Innate Lymphoid Cells by IL-33 to Prevent Renal Ischemia-Reperfusion Injury. J Am Soc Nephrol 2018;29:961–76.

30. Chen F, Yang W, Huang X, et al. Neutrophils Promote Amphiregulin Production in Intestinal Epithelial Cells through TGF-beta and Contribute to Intestinal Homeostasis. J Immunol 2018;201:2492–501.

31. Zhang J, Ji JY, Yu M, et al. YAP-dependent induction of amphiregulin identifies a noncell-autonomous component of the Hippo pathway. Nat Cell Biol 2009;11:1444–50.

32. Lefort S, Tan S, Balani S, et al. Initiation of human mammary cell tumorigenesis by mutant KRAS requires YAP inactivation. Oncogene 2020;39:1957–68.

33. Kobayashi S, Nagafuchi Y, Okubo M, et al. Integrated bulk and single-cell RNA-sequencing identified disease-relevant monocytes and a gene network module underlying systemic sclerosis. J Autoimmun 2021;116:102547.

34. Gao X, Jia G, Guttman A, et al. Osteopontin Links Myeloid Activation and Disease Progression in Systemic Sclerosis. Cell Rep Med 2020;1:100140.

35. Casasanta MA, Yoo CC, Udayasuryan B, et al. Fusobacterium nucleatum host-cell binding and invasion induces IL-8 and CXCL1 secretion that drives colorectal cancer cell migration. Sci Signal 2020;13.

36. Amorim NRT, Souza-Almeida G, Luna-Gomes T, et al. Leptin Elicits In Vivo Eosinophil Migration and Activation: Key Role of Mast Cell-Derived PGD2. Front Endocrinol (Lausanne) 2020; 11:572113.

37. Yeh TC, Marsh V, Bernat BA, et al. Biological characterization of ARRY-142886 (AZD6244), a potent, highly selective mitogen-activated protein kinase kinase 1/2 inhibitor. Clin Cancer Res 2007;13:1576–83.

38. Blyszczuk P, Kozlova A, Guo Z, Kania G, Distler O. Experimental Mouse Model of Bleomycin-Induced Skin Fibrosis. Curr Protoc Immunol 2019;126:e88.

39. Fan Q, Gu D, Liu H, et al. Defective TGF-beta signaling in bone marrow-derived cells prevents hedgehog-induced skin tumors. Cancer Res 2014;74:471–83.

40. Shao X, Liao J, Lu X, Xue R, Ai N, Fan X. scCATCH: Automatic Annotation on Cell Types of Clusters from Single-Cell RNA Sequencing Data. iScience 2020;23:100882.

41. Gu D, Lin H, Zhang X, et al. Simultaneous Inhibition of MEK and Hh Signaling Reduces Pancreatic Cancer Metastasis. Cancers (Basel) 2018;10.

