## Supplementary material for "A critical role of AREG for bleomycin-induced skin fibrosis": Figure on single cell RNA-Seq

A

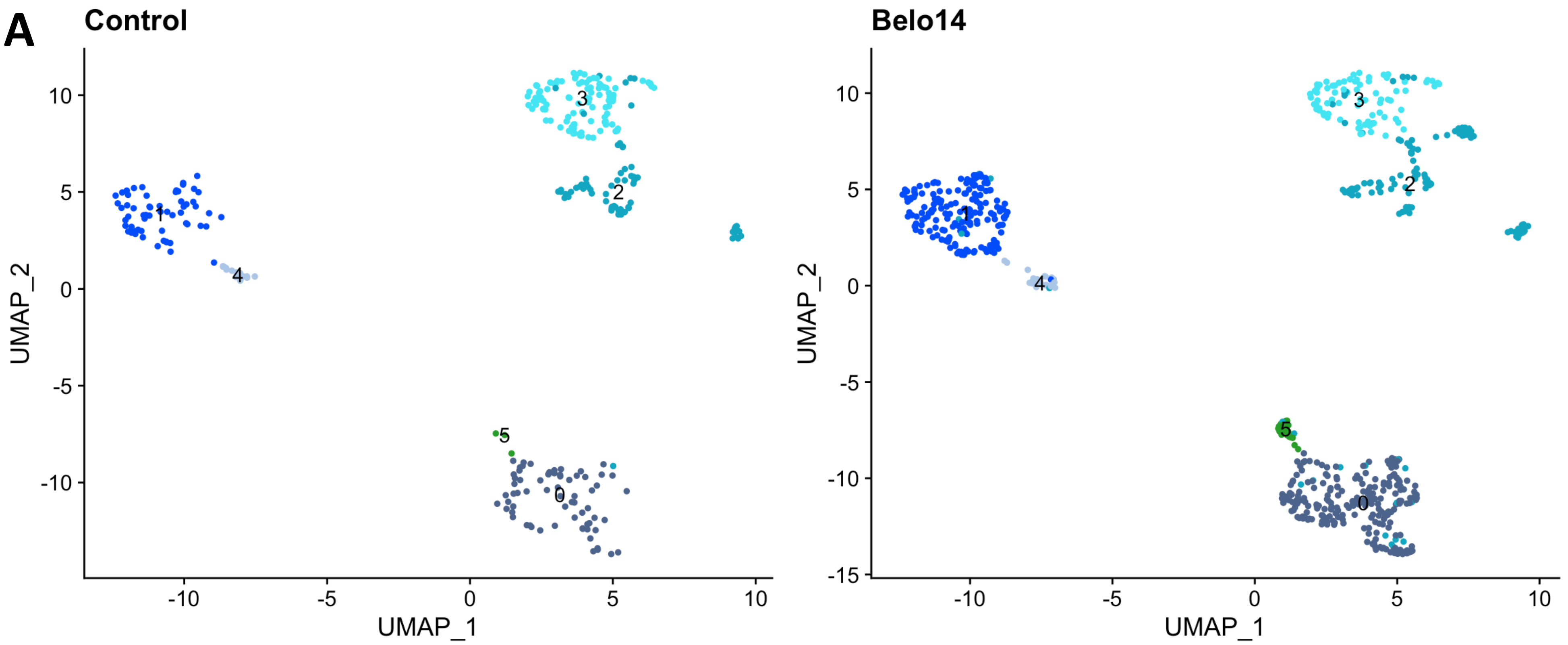


B

| Areg expression in clusters | Cluster | 0 | 1 | 2 | 3 | 4 | 5 |
|---|---|---|---|---|---|---|---|
| Control | | - | + | + | + | + | + |
| Bleomycin-injected (Bleo14) | | - | + | + | + | - | - |

**Supplementary data 3- A summary of single cell RNAseq analyses and *Areg* expression in different cell populations. A** shows distribution of cell clusters in control and bleomycin-treated mouse skin. **B** summarizes *Areg* expression in different cell clusters. Cluster 0 contains macrophages and monocytes. Cluster 1 contains keratinocytes. Cluster 2 contains immune cells and endothelial cells. Cluster 3 contains fibroblasts (and myofibroblasts). Cluster 4 contains NK cells and T cells. Cluster 5 contains dendritic cells.
